# Longitudinal single cell RNA-sequencing reveals evolution of micro- and macro-states in chronic myeloid leukemia

**DOI:** 10.1101/2025.05.14.653262

**Authors:** David E. Frankhouser, Dandan Zhao, Yu-Hsuan Fu, Anupam Dey, Ziang Chen, Jihyun Irizarry, Jennifer Rangel Ambriz, Tiffany Kanesa Ybarra, Sergio Branciamore, Denis O’Meally, Ryan S. Sathianathen, Jeffery M. Trent, Stephen Forman, Adam L. MacLean, Ya-Huei Kuo, Kathleen M. Sakamoto, Bin Zhang, Russell C. Rockne, Guido Marcucci

## Abstract

Single cell RNA sequencing (scRNA-seq) has revolutionized our understanding of cancer, yet identifying meaningful disease states from single cell data remains challenging. Here, we systematically explore the chronic myeloid leukemia (CML) specific information content encoded in single cell versus bulk transcriptomics to resolve this paradox and clarify how discrete disease-defining states emerge from inherently noisy single cell data. We demonstrate that, while CML single cell transcriptomes exist along continuous transcriptional microstates, clinically relevant leukemia phenotypes clearly manifest only at the pseudobulk (macrostate) level. By leveraging state-transition theory, we reveal how robust disease phenotype state-transitions are governed by cell type specific contributions. Our results establish a theoretical framework explaining why discrete disease phenotypes remain hidden at the single cell scale but emerge clearly at the aggregated macrostate level, enabling previously inaccessible biological insights into leukemia evolution. By resolving how single-cell variation aggregates into macroscopic disease states, our framework provides new insight into CML progression and offers a broadly applicable strategy for exploring disease dynamics across cancers and other complex conditions.

## INTRODUCTION

Single cell RNA sequencing (scRNA-seq) provides unprecedented detail into cellular states and heterogeneity in cancer yet translating this detailed information into clinically meaningful chronic phase (CP) and blast crisis (BC) disease states remains challenging. A fundamental but unresolved question is how the continuous, granular information at the single cell level relates to the discrete disease phenotypes readily observed using bulk transcriptomics. While bulk data effectively captures robust disease states, single cell analyses often yield a complex continuum of transcriptional states that may obscure clear phenotype boundaries and complicate biological interpretation and clinical relevance.

To bridge this gap, we explore the information content of single cell compared with bulk-level transcriptomic data. We introduce a conceptual framework grounded in state-transition theory to explain an apparent paradox that emerged from our data: although individual cells exhibit continuous transcriptional variation at the single cell level (microstates), clinical phenotypes emerge clearly only at the aggregate or pseudobulk (macrostate) level. We apply our theoretical approach to chronic myeloid leukemia (CML), a disease defined by the presence of the BCR::ABL fusion gene and characterized clinically by two distinct phases: initial CP and later BC. Despite successful achievement of long-term remission with tyrosine kinase inhibitors, the persistence of leukemia stem cells in CP, that could eventually drive disease transformation into BC, highlights the need for better understanding of leukemia dynamics and makes CML an ideal model system to explore how information of single cells relates to disease phenotype dynamics. To date, CML studies using scRNA-seq have primarily been used to either investigate stem and progenitor cell populations^1–3^ or to identify how specific immune cells respond or influence therapy^4,5^. Here, we approach CML at the system-level, and using state-transition modeling of scRNA-seq data, we quantify how the population and transcriptional changes of all peripheral blood cell types contribute information to CML development and dynamics.

## RESULTS

### CML state-transition is not detectable at the single cell level

Our previous state-transition work constructed a CML state-space from time-series bulk RNA-seq (transcriptome) data^6^. Using principal component analysis (PCA) by performing singular value decomposition (SVD) on the time-series gene expression matrix, we showed that disease trajectories were encoded in the CML state-space corresponding to the first principal component (PC1)^6,7^. The state-space comprised a continuum of transcriptomic states that spanned from health to disease.

To test how a state-transition model could also be derived and whether it could be more informative using transcriptomes of individual cells, we conducted a time-series single cell RNA-seq analysis on peripheral blood mononuclear cells (PBMC) samples collected over time from a BCR::ABL inducible mouse model of CP CML. Mouse blood was sampled before induction of BCR::ABL by tetracycline withdrawal at week 0 (T_0_) and then weekly for 10 weeks after induction (11 samples per mouse; 29 total samples; Fig 1A). Because the mice developed CML at different rates, we scaled the time points to represent a progression from the first time (T_0_) to the last collection time point (T_f_), which was either before a moribund mouse was culled or end of the study (week 10), whichever occurred first. We used the scaled time to identify whether a candidate state-space ordered the mouse time point samples from healthy time point (T_0_) to the leukemic time points (T_f_). To construct a single cell leukemia (or CML) state-space, we applied SVD to the sc gene expression matrix and visualized scaled time in each resulting PC (Fig. 1A). One of the BCR::ABL-induced mice did not develop leukemia over time and therefore, did not have a leukemic time point (T_f_). We observed that the first two PCs grouped cells by cell type rather than by disease status, and, in fact, none of the first 6 PCs or UMAP analysis could separate cells by disease state (i.e., health vs leukemia; Fig. S1B-C). We therefore concluded that the difference in cell types, rather than in disease status, was the primary source of variance in the sc transcriptomes. Thus, we examined whether leukemia state-space could be identified by analyzing each cell type independently. To this end, we categorized the PBMCs into four main cell types: B-, T-(including NK-cells), Myeloid, and Stem cells based on cell type annotation analysis (Fig. 1A; Table S1). When SVD was performed on the sc gene expression matrix of each of the four cell types individually, no PC for any of the cell types produced a state-space that separated health from leukemia; in fact, T_f_ cells were never associated with a unique leukemic transcriptional state (Fig. 1B; S1D). Other methods for scRNA-seq analysis^8,9^ were also unable to identify a state-transition from heath to leukemia at the sc-level (Fig. S1E-F). Of note, despite not observing a unique leukemic state-space at the sc-level, we observed differentially expressed genes (DEGs) in each cell type when gene expression at T_0_ was compared with that at T_f_ indicating that individual gene expression does change during leukemia progression, but not sufficiently to overcome the cell type signature or to identify the system phenotype transition in the sc transcriptomes (Fig. S1G).

**Figure 1:**
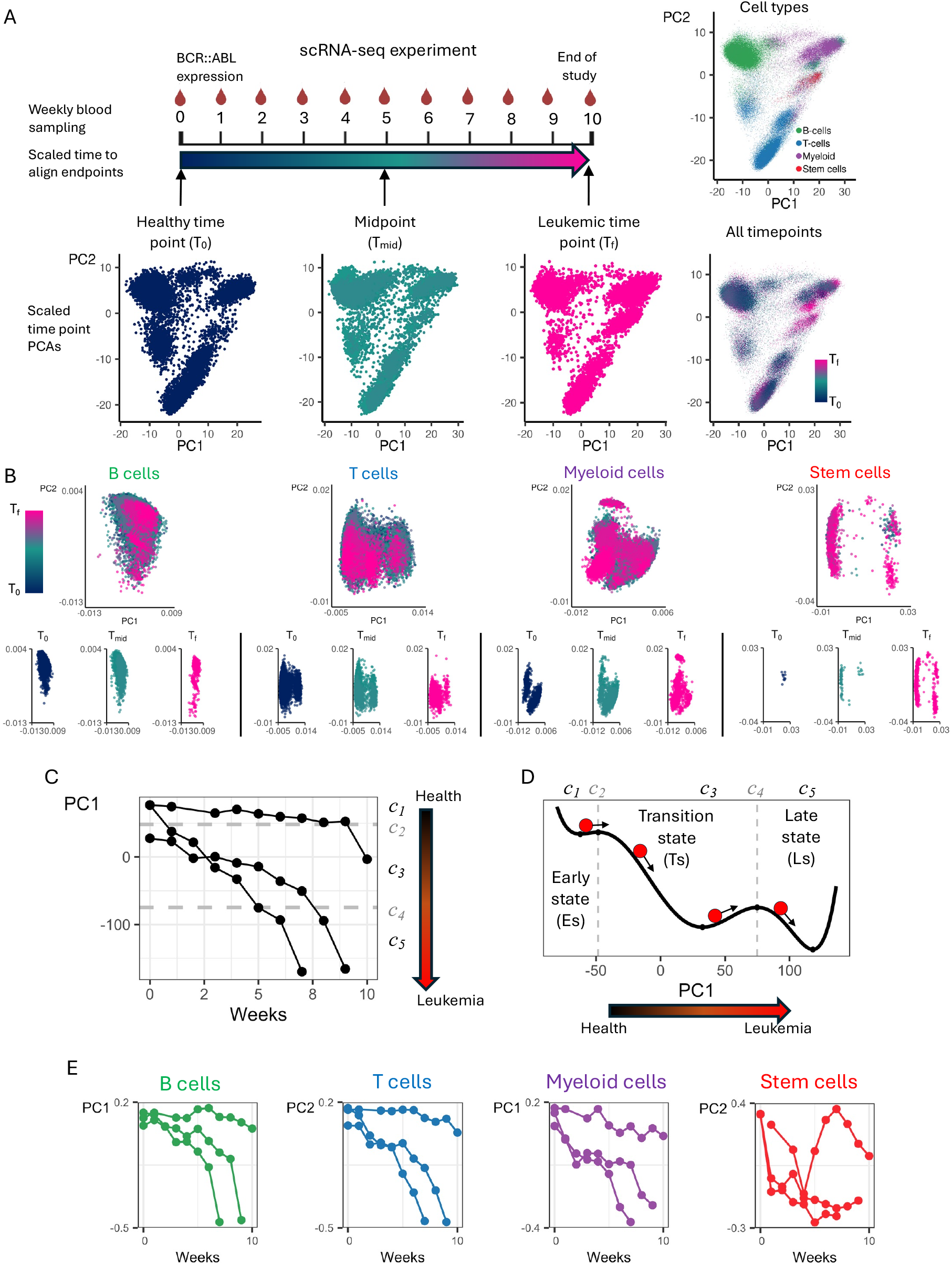
The leukemia phenotype is a macrostate observable in the pseudobulk data. **A**) Time series PBMC samples from a chronic phase CML mouse model were collected weekly and scRNA-seq was performed to determine the cell contribution to leukemia state-transition (top left). Because the mice developed leukemia at different rates, a scaled time was used to align the samples from the healthy (T_0_) to leukemia (T_f_) time points. PCA was performed using all time points and all cells to determine if a unique transcriptional state associated with leukemia (T_f_) could be identified (i.e., a sc-level CML state-space; bottom). After labeling cells by cell type and visualizing cell type in PCA space, the primary source of variance (PC1 and PC2) separated cells by cell type and not leukemic from healthy time points. **B**) PCA was performed on each cell type separately to determine whether a CML state-space could be identified within each cell type. In all PCs, the cells were observed to be in a continuum of transcriptional states that included states from both health and leukemic time points. **C**) Pseudobulk transcriptomes for each sample were created using all cells and, following the same procedure used previously to identify a CML state-space using bulk RNA-seq, a PsB CML state-space was identified in PC1. When plotted vs time, each mouse’s trajectory in PC1 shows the CML development as the mice move from the healthy to leukemic state. The sample density in the state-space was used to identify boundaries between stable critical points (*c*_1_, *c*_3_, *c*_5_) and the unstable boundary critical points (*c*_2_, *c*_4_). **D**) Using the critical points of the PsB state-space, a potential landscape was constructed that describes the sample’s CML dynamics. Each mouse is modeled as a particle undergoing Brownian motion in the potential landscape. The stable critical points are used to define the Early, Transition, and Late phenotypic states of CML. **E**) Cell type PsB transcriptomes were created for each sample by combining gene counts from the cells of each cell type. Using the same procedure to identify the PsB state-space, a CML state-space was identified in either PC1 or PC2 for each cell type. The ctPsB state-spaces showed different leukemia development dynamics when compared to the PsB state-space.

Taken together, these results suggest that, from T_0_ to T_f_, the cells occupied a continuum of transcriptional states that always overlapped healthy and leukemia state (Fig. 1B; Fig. S1B-F). We describe this continuum of transcriptional states in each lower dimensional representation of the data as a superposition of cell states.

### CML disease evolution is observable in the pseudobulk

As no single leukemic state emerged from the sc-level transcriptome analysis, we then asked whether the individual cells when aggregated can define the disease phenotype. To prove if disease phenotype dynamics (e.g. health to leukemia transition) encoded by the cell microstates could be identified when we considered these microstates in aggregate (i.e., macrostates), we constructed pseudobulk (PsB) transcriptome samples by combining gene counts from all individual cells. After performing SVD on the PsB gene expression matrix, we observed a trajectory of the PsB transcriptomes from healthy state (T_0_) to leukemic state (T_f_) in PC1. This trajectory was also the only PC that correlated with BCR::ABL expression levels (Fig. S1H; Table S2) and therefore identifiable as the PsB CML state-space (Fig. 1C). We then aligned the PsB CML state-space and a bulk CML state-space that we previously reported^6^, and plotted them together over time to compare how their respective disease dynamics mapped in the state-space (Fig. S1K). Like in the bulk state-space^6^, the disease dynamics in the PsB state-space was described by a leukemogenic potential with three steady states (or “wells”) (Fig. 1D). The three wells of the leukemogenic potential were used to define three disease states of leukemia (stable critical points) and we labeled them as follows: Early state (Es) at *c*_1_, Transition state (Ts) at *c*_3_, and Late state (Ls) at *c*_5_. The three-well potential also had two unstable critical points which served as the boundaries between the steady states and that we labeled as: Early-Transition state (T-Es) at *c*_2_, and Transition-Late state (T-Ls) at *c*_4_.

### Each cell-type undergoes CML state-transitions

Based on the observation that the CML state-transition was encoded by the total PsB data, we then tested if we could identify a state-space that encoded the transition from health to leukemia also using individual cell type PsB transcriptomes. To this end, we created PsB transcriptome samples for each of the four PBMC cell types (ctPsB) and performed SVD separately on each ctPsB gene expression matrix (Fig. S2A). Using the scaled time ordering of the samples and BCR::ABL expression for each of the four cell types, we were able to identify the CML state-space in either the first or second PC for each of the four cell types (Fig. 1E). Like the PsB state-space, the B-, T-, and Myeloid ctPsB all had tristable potentials (Fig. S2B). In contrast, likely due to the smaller number of cells in each sample, the trajectories for the Stem cells were noisier and appeared to exhibit bistable dynamics (Fig. S2B). This result demonstrated that individual cells indeed contained information about disease state-transition that can be extracted only when the sc-level microstates are considered in aggregate using PsB transcriptomes.

To explore how each ctPsB contributed to the total PsB state-space, we compared the trajectory of the total PsB transcriptomes with those of individual ctPsB transcriptomes mapped into the total PsB state-space (Fig. S2C). We noted that each projected ctPsB trajectory started and ended at different locations of the state-space. To summarize the ctPsB projections, we defined the span of their trajectories to be the minimum and maximum observed state-space coordinate for each cell type (Fig. S2D). We noted that the span of each cell type only occupied a subset of the PsB trajectories, and that each span had different lengths. Although none of individual ctPsB trajectories fully spanned the total PsB state space, when considered together, they completely overlapped the total PsB state-space, suggesting that the latter is a linear combination of the former and that each cell type contributed differently to the disease dynamics.

### State-transition reveals cell type-specific contributions to CML development

To understand the contribution of individual cell types to disease dynamics, we then plotted the proportions of each cell type over the scaled time for each leukemic mouse to visualize the changes that occurred during CML development (Fig. 2A). To this end, we grouped the samples into the three disease states (Es, Ts, Ls) using their state-space coordinates and compared the cell type proportions of the individual cell types at each disease state with those observed at the healthy (T_0_) state (Table S3; Fig. 2A). The Stem cells were the first to show significant population changes that started at Ts (increased; p < 0.001) and continued to increase to Ls (p=0.001). The Myeloid and B cells did not change until Ls with the Myeloid cell population increasing (p = 0.03) and B cell population decreasing (p = 0.03). The T/NK cell population size did not show significant changes.

**Figure 2:**
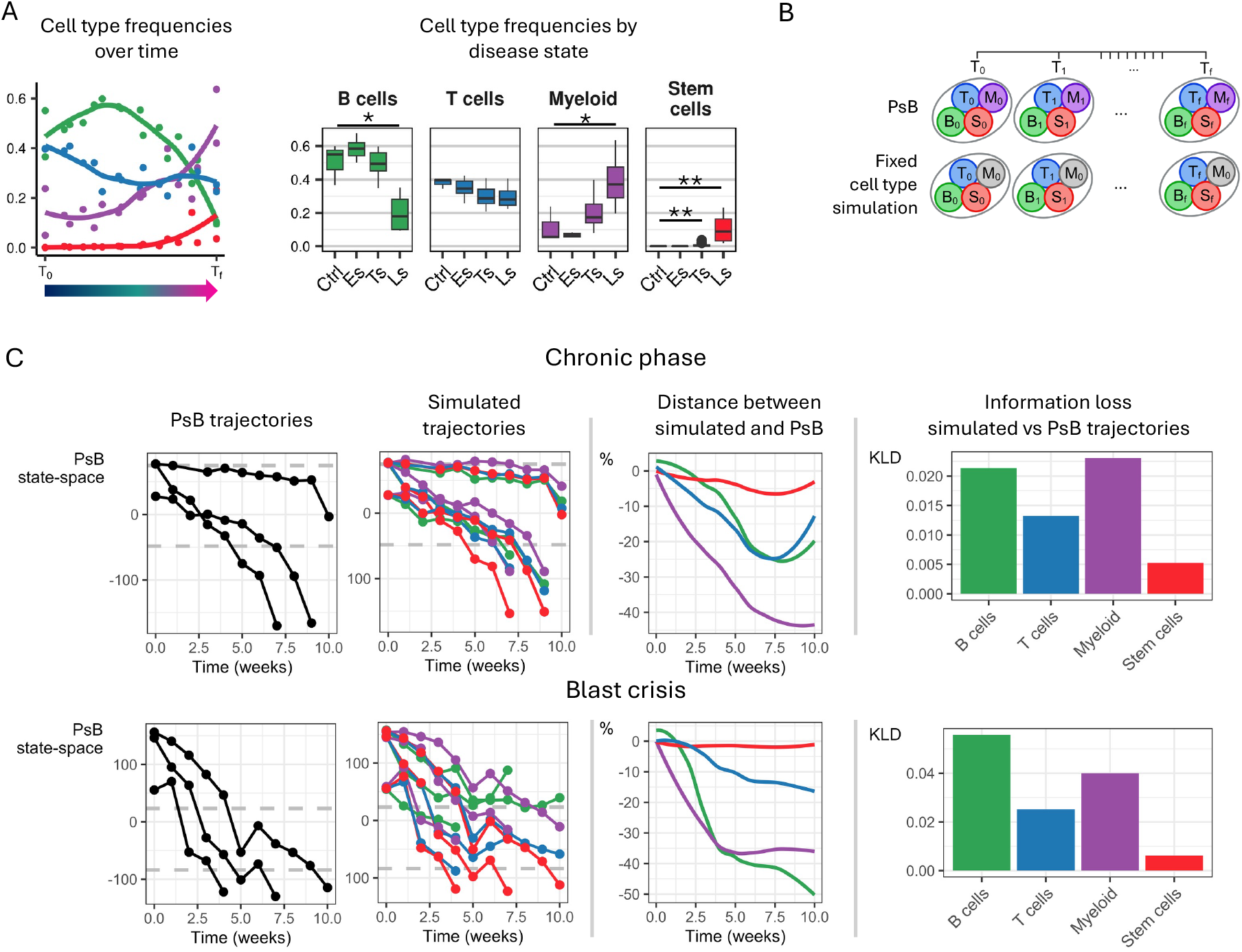
Cell types contribute different information to leukemia dynamics. **A**) Number of cells in each cell type are shown as a proportion over the scaled time of leukemia development (left). The cell type proportions are grouped by disease state to test cell type population changes over leukemia development (* p < 0.05; ** p ≤0.001). **B**) The computational simulation strategy to quantify each cell type’s contribution to CML dynamics is depicted in the diagram. Each cell type was fixed at the healthy (T_0_) time point by replacing all cells of that type with the healthy time point cells for all subsequent time points. Those cells were then used to construct a simulated PsB sample for each time point sample to simulate the effect of each cell type being fixed at T_0_ cell population count and gene expression state. **C**) (top) The full PsB trajectories for the CP CML mice were compared to the simulated trajectories where each cell type is fixed at the healthy (T_0_) time point (left). For each sample, the distance between the full and simulated trajectories was calculated as a percentage and then summarized by each cell type to show how much less state distance the fixed cell type simulations traveled in the state-space (middle). To quantify the information loss from fixing each cell type, Kullback–Leibler divergence (KLD) was calculated between the full PsB and simulated trajectories for each cell type to show the Myeloid cells contributed the most information to the state-transition if not allowed to evolve. (bottom) The same simulations were performed for the BC CML mice which showed different contribution from the cell types with B cells contributing the most information to BC CML state-transition.

Next, we performed a computational simulation that leveraged both the time-series samples and the PsB state-space to quantify the contribution of each cell type to CML development. To simulate the effect of not having one of the cell types evolve over disease development, we computationally subtracted the contribution of each cell type by fixing both the cell counts and gene expression of each of the cell types individually to the values observed at the healthy state (T_0_) (Fig. 2B). We projected all time points of the “fixed cell type” data into the space to construct simulated trajectories of the “fixed cell type” PsB transcriptomes. We then compared each “fixed cell type” simulated trajectory to the experimentally derived (observed) PsB transcriptome trajectories (Fig. 2C). To quantify the contribution of each cell type to the state-transition from health to leukemia, we calculated the difference between the simulated and observed trajectories at each time point (Fig. 2C middle). We postulated that if a fixed cell type did not shorten the state-space trajectories compared to the observed PsB trajectories, then that fixed cell type likely did not contribute significantly to the PsB state-transition from health to leukemia. Conversely, we interpreted a larger the distance between observed and simulated trajectories as lost information due to the lost contribution of the fixed cell type. To quantify information loss, we calculated the Kullback-Leibler divergence between the observed and the “fixed cell type” simulated PsB trajectories for each cell type (Fig. 2C right). We observed that fixing the Myeloid cells was associated with the largest information loss, suggesting that this cell population contributed the most to CML state-transition. This result was also supported by the observation that Myeloid cells also had the largest span between the projected ctPsB and PsB trajectories in the state-space (Fig. S2D) and therefore the largest contribution to CML state-transition which is likely due to the combination of the progressively increasing cell number (Fig. 2A) and the gene expression changes (Fig. S1G) over time. Although the myeloid cells contributed the most information, we noted that all cell types contributed information to the disease phenotype which further supports our macroscopic view of disease.

### An independent CML model confirms the state-space is encoded in PsB data

To confirm our findings, we conducted a second time-series scRNA-seq experiment using a different mouse model that recapitulates blast crisis CML development^10^. While the CP CML mouse was a solely BCR::ABL inducible model, the BC mouse, in addition to inducible BCR::ABL expression, had a germline miR-142 knock-out as a “second leukemogenic hit” that triggered transformation. While genotypic differences between these two models prevented direct comparison of the transcriptional and disease states, we used the same analytic approach to identify a sc-level state-space for the BC model. Like the CP model, we were unable to identify state-transition encoded in the sc-level transcriptomes, but we could identify a leukemic state-space in PC1 when sc information was aggregated into the PsB transcriptomes (Fig. S3A-C). Based on the BC CML PsB state-space trajectories, we observed that BC CML exhibited bistable phenotypic dynamics (Fig. S3D) that were described by a two-well potential labeled as early and late disease states (Fig S3D). When we constructed ctPsB for the BC CML samples, we observed that each cell type encoded state-transition and spanned different portions of the PsB state-space (Fig. S3E-G). Although BC CML has different dynamics when compared to CP CML, the BC CML experiment validated our main findings that the leukemic state-transition is not encoded at the sc-microstate level but is observed only when PsB macrostates are considered.

Finally, we applied the fixed cell type simulation strategy to the BC CML samples to quantify the contribution of each cell type to BC CML PsB state-space (Fig. 2C bottom) using Kullback-Leibler divergence to assess the difference in information between the simulated vs PsB trajectories. In the BC CML model, we observed that B cells and Myeloid were the cell types that contributed the most information to state transition (Fig. 2C right).

### Validation of the macroscopic state-transition in human CML

Our experimental setup has two potential confounding factors: 1) in our mouse model, BCR::ABL is induced in every cell type which could homogenize the cell state changes, and 2) profiling may under-represent the transformed progenitor/stem compartments that define disease. To address both in the case of natural progression, we analyzed CD34^+^ selected bone marrow (BM) scRNA-seq from CML patients in chronic phase at diagnosis and age-matched healthy controls (n=4 CML, n=3 healthy). As in the mouse experiments, PCA and other dimensionality reduction methods at the single cell level did not reveal a disease specific transcriptional state. When all cells were analyzed together, PCA and UMAP both separated cells by lineage rather than disease state (Fig. 3A; S4A-B). The same was true when we analyzed each cell type independently which showed that in the top 5 PCs, the healthy and CML cells occupied overlapping, continuous transcriptional states (Fig 3B; S4C). Thus, we concluded that disease status was not encoded as discrete single cell states in the human BM data. In contrast, when we aggregated cell counts into PsB samples, the disease macrostate emerged as a unique transcriptional state. PC2 of the PsB PCA identified a state-space that could cleanly separate CML from healthy samples (Fig. 3C). Together these results reproduce what we observed in the mouse models: the cell level data constitutes microstates of the disease phenotype macrostate.

**Figure 3:**
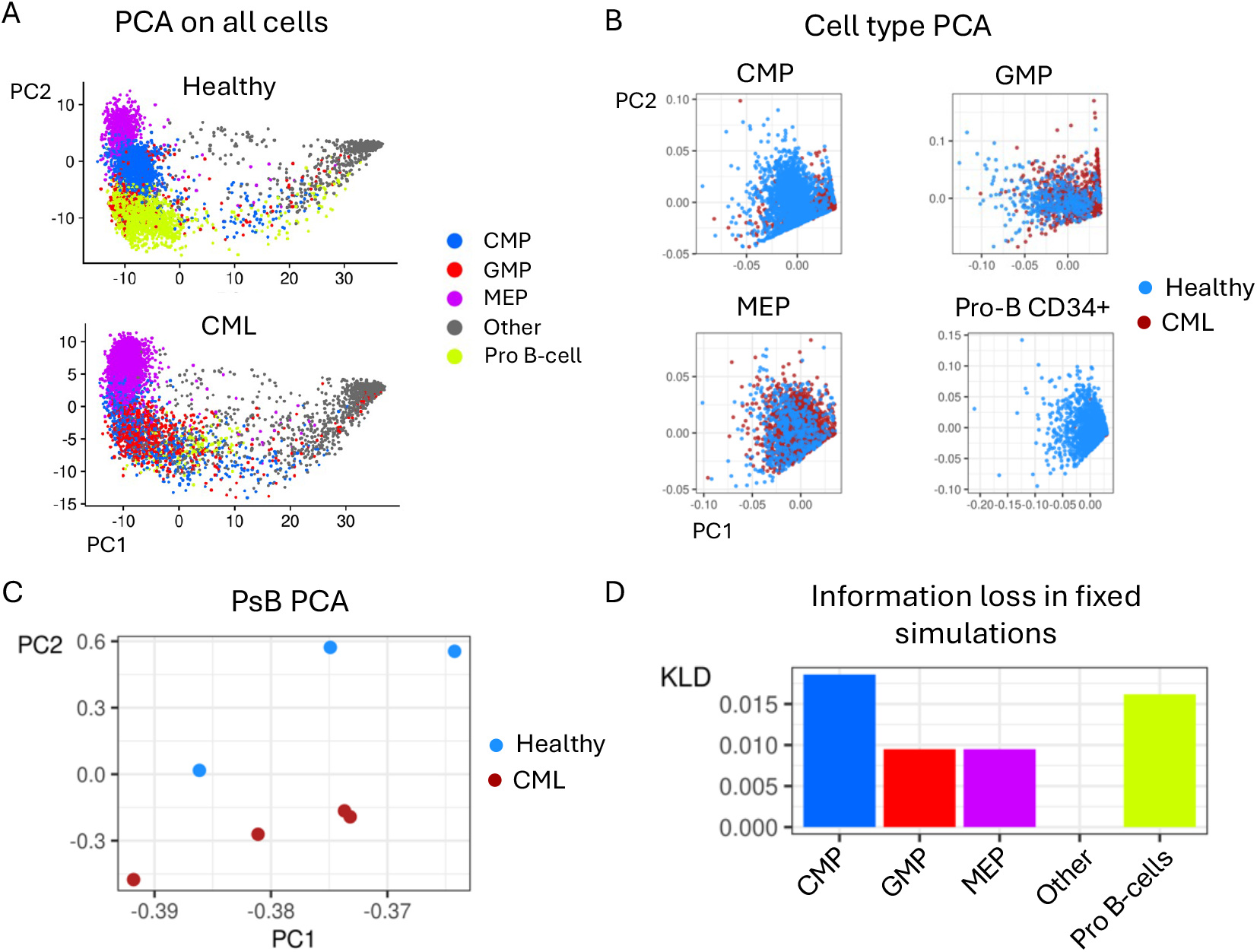
Validation on human CD34+ BM cells. **A**) Using a cohort of 4 CML and 3 healthy BM samples, CD34^+^ cells were isolated and scRNA-seq was performed. Cell types were labeled from the data and PCA was performed to determine whether a unique transcription disease could be identified from the data. However, similar to the mouse model, cell type and not the disease status of the samples was the primary source of variance. **B**) PCA was also performed on each cell type separately to identify whether disease state emerged at the single cell level, but the healthy and CML cells were observed to be in a continuum of transcriptional states in all PCs. **C**) The fixed simulations were used on all pairwise combinations of the healthy and CML samples to quantify how each cell type contributed to CML. For each pairwise healthy-CML sample pairing, the Kullback–Leibler divergence (KLD) was calculated to quantify how much information was lost for each cell type and the mean from all pairwise pairings was reported. The common myeloid progenitors (CMP) contributed the most information whereas the “other” cell type group that combined all remaining cell types accounting for less than 5% of the total cells in both healthy and CML samples contributed almost no information to CML.

Because these data are from different individuals, we adapted our fixed cell-type simulation to quantify contributions without matched time-series samples. We used all healthy–CML combinations, and for each pairing and each cell type, we “fixed” that cell type by replacing its CML cells with the paired healthy cells (Fig. S4D). We then recomputed the PsB state-space for the fixed simulations on each cell type (Fig. 3D *left*) and measured the shortfall in the state-space distance relative to each healthy-CML difference (Fig. 3D *middle*; Fig. S4D). The information lost from all fixed simulation combinations was calculated using the Hullback-Leibler divergence to show that common myeloid progenitors (CMP) and Pro B-cells contributed the largest share of information to the human CML, whereas the pooled “other” category contributed negligible information (Fig. 3D). Similar to the mouse results, the fixed simulation on human BM data confirms that multiple cell types contribute information to the CML macrostate.

### Cell type contributions are modeled as linear combinations of their transcriptomic state

We showed that each of the PBMC cell types contribute in different proportions to the CML state-transition. To model the dynamics of state-transition mathematically, we postulated that the total BCR::ABL signal from the PsB (*C*_*PsB*_) acts on each cell type and affects their ctPsB transcriptome (Fig. 4A). A model that incorporates a linear combination of individual cell types to rederive the PsB dynamics could be expressed as 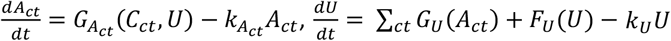 where *A*_*ct*_ represents the four cell type populations, *U* is the PsB transcriptome^11^. The functions 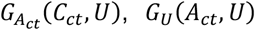 and *F*_*U*_(*U*) are composed of Hill functions of the form 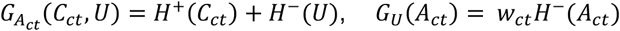 and *F*_*U*_(*U*) = *H*^+^(*U*), where *H*^+^ represents a Hill function for activation and *H*^−^ represents a Hill function for repression (see methods (model description in blue text)). *w*_*ct*_ is a weight factor whose value governs the contribution of each cell type to the PsB. The potential of the system^11^ is the integral of the leukemic “force” which depends on the BCR::ABL signal, so that:

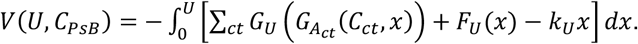

**Figure 4:**
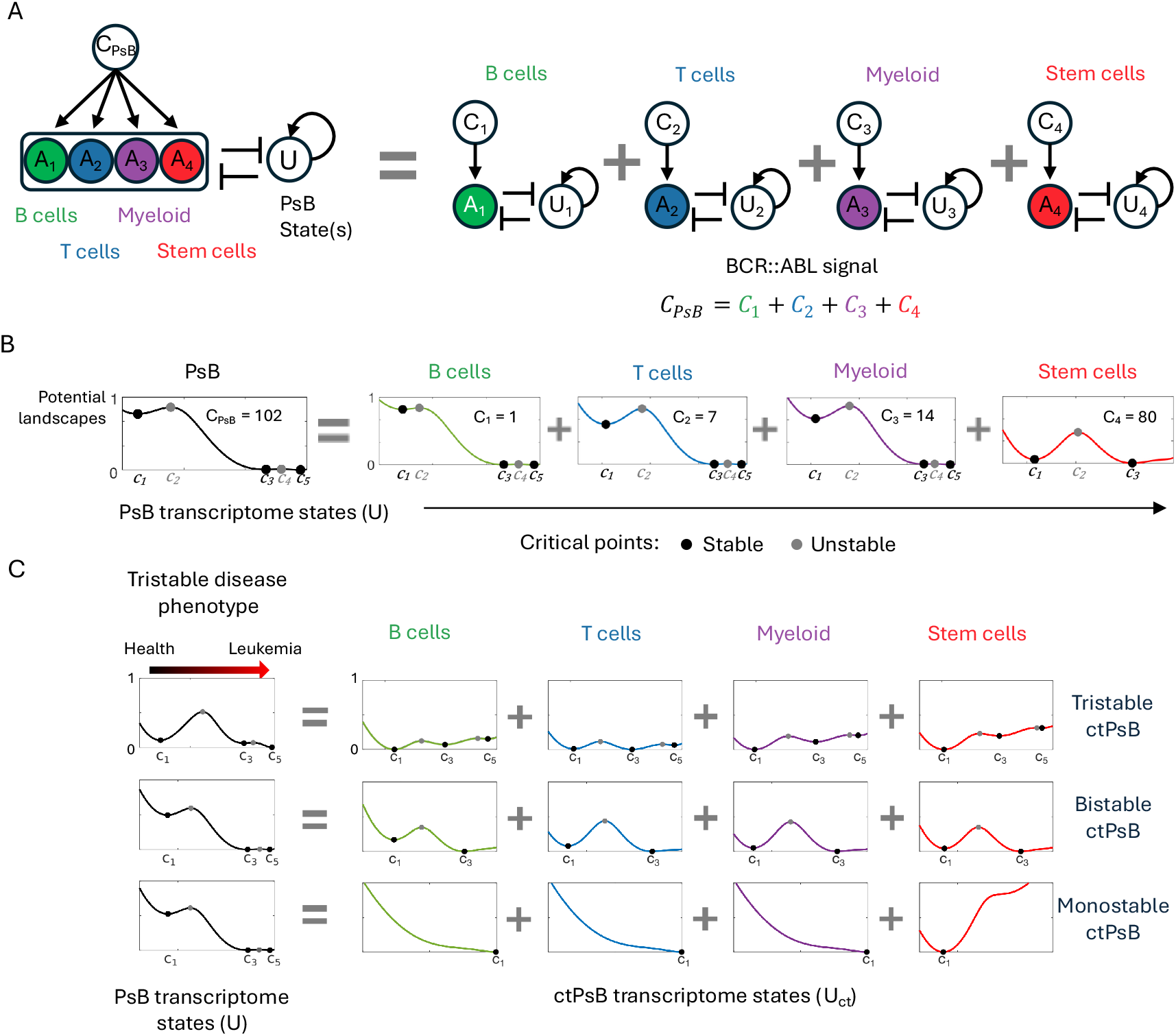
Math model relates cell type dynamics to the disease phenotype. **A**) Cell type PsB state-space dynamics were incorporated into a mechanistic model that related the total BCR::ABL CML oncogene signal computed from the pseudobulk (*C*_*PsB*_) to the PBMC population cell types (*A*_*ct*_) and the PsB transcriptome state-space (*U*). The PsB state-space was reconstructed by adding each ctPsB model together as linear combination of each ctPsB model. The ctPsB models had cell type specific parameters including the BCR::ABL signal (*C*_*ct*_) which was based on the observed BCR::ABL expression in each cell type. The total BCR::ABL signal in the PsB (*C*_*PsB*_) was the sum of the signals from each cell type. **B**) BCR::ABL expression in each cell type and ctPsB state-space dynamics observed from the data informed the parameter used to construct a CP CML model. In this model, the cell type dynamics are combined to reconstruct the three-well potential that describes the PsB state-space phenotype dynamics. **C**) The mechanistic model demonstrates how different cell type dynamics can combine to produce tristable dynamics in the PsB phenotype. Even simple monostable cell types can produce a tristable phenotypic dynamics.

To apply this model to our CP CML scRNA-seq data, we used both the ctPsB state-space dynamics (Fig 1A) and the BCR::ABL expression in each cell type (Fig. S2E) to determine the number and location of the steady states for each cell type. From the critical points, we constructed the potentials that define state-transition dynamics for each cell type (Fig. 4B). We modeled the B-, T-, and Myeloid cells as having tristable dynamics, and Stem cells as bistable. Individual cell type models were constructed by setting the two parameters that controlled the cell type dynamics so that each cell type model matched the observed ctPsB state-space and projected dynamics (Fig. 4B; Table S4). The result of this model recapitulated a tristable CML potential constructed from the cell type transcriptional dynamics, suggesting that the disease phenotype dynamics is a linear combination of the distinct ctPsB from health to a leukemic state.

While fitting the model to the data, we noted that the tristable potential observed in the system-level PsB state-space could be produced using combinations of different cell type dynamics including cell type potentials that were all monostable, bistable, or tristable (Fig 4C). This result demonstrated how even monostable dynamics in cell populations can combine to produce tristable dynamics as observed in the data. Here, the theory reinforces our understanding of the information hierarchy embedded in disease state-transitions. The information from cell level microstates can be aggregated into cell type population dynamics (ctPsB). While certain cell types exhibited tri- and bistable states, cell type dynamics could also manifest as simpler monostable states, where cells transition from their original stable state to occupy a single new state. These dynamics can again be aggregated using the mechanistic model of disease to finally produce system level phenotype dynamics (CML state-space), which describes the disease trajectories of individuals. This mechanistic model demonstrates how the distinct dynamics of blood cell populations integrate to produce the overall leukemic trajectory, providing a quantitative framework that could ultimately be applied to patient samples to predict disease progression, stratify risk of transformation, and guide therapeutic decision-making.

## DISCUSSION

Here, we studied the evolution of CML from the point of view of an epigenetic state-transition of the peripheral blood transcriptome. While it is widely recognized that cells undergo state-transitions during normal and pathologic differentiation^12–19^, we studied this process with longitudinal sequencing data from a mouse model of CML with the goal of understanding the relationship between single cell state-transitions and phenotype transition from health to disease. Single cell RNA sequencing data presents significant analytical challenges, including limited read depth, dropout events, and complexities in interpreting longitudinal dynamics^20–26^. For these reasons, we guided our experimental and analytical approach with a theoretical and mathematical framework^6,7,27^. We first showed that CML transcriptional state-transition was not encoded in the cell level data, where cells were found to exist in a continuum of transcriptional microstates in a superposition of health to disease states. Instead, the phenotype state-transition became apparent only after information derived from single cell microstates was integrated into macrostates by constructing PsB transcriptomes.

To explore state-transitions in different cell populations, we performed PsB for each cell type (ctPsB) and showed that each cell type had both a unique state-space and distinct CML state-transition dynamics. We developed a computational simulation to quantify each cell type’s contribution to CML by fixing each cell type population at its healthy state for all subsequent time point samples and determined information loss by comparing each fixed cell type simulation to the experimentally observed PsB trajectories. In our CP mouse model, we identified Myeloid cells as the cell type that contribute the largest amount of information to the system dynamics. While this is not surprising given that we used a CML mouse model, it is important to underscore that this conclusion was derived experimentally from an unbiased data analysis, thereby validating the usefulness of this approach to CML and its potential applicability to other types of leukemia and cancer to predict disease evolution or treatment response^6^. To this end, these state-transition results were validated in a second time-series experiment on a BC CML model and found to be consistent when CD34^+^ selected cells from human BM samples were analyzed. Despite differences in genotype and CML disease model, both experiments demonstrated that the sc level data encoded a microstate of the PsB phenotype dynamics. Finally, we derived a mechanistic model where the PsB transcriptome dynamics can be recapitulated as a linear combination of the ctPsB dynamics and the cell type-dependent effects such as BCR::ABL expression. The mechanistic model provided a formal framework to understand how cell types with different transcriptional dynamics can produce complex system-level phenotype dynamics.

Taken together, we illustrate here how disease-relevant information is contained from data collected at different biological scales. We used the state-transition model to discover a relationship between sc and PsB/system-level data. We described this relationship using the thermodynamic concepts of micro- and macrostates as previously hypothesized^28–30^, but not yet experimentally demonstrated through longitudinal single cell analysis^31^. At the sc-level, cells exist in a continuum of microstates that include both healthy and disease transcriptional states. While we were unable to recover disease state transition from sc microstates, the system phenotype (i.e., disease transition) can be recovered at the macrostate level when sc microstates are considered in aggregate (i.e., PsB). Thus, the macrostate may be considered as a continuous variable that describes phenotype evolution within a state-space that ranges from healthy to disease. For both CP and BC CML models, we showed that macrostate could be identified using either the PsB data from all PBMCs or using the ctPsB data from any one of the cell types that comprise the PBMCs. Given a PsB transcriptome from any of these sources, we also discovered that the disease macrostate emerged from aggregated sc microstates, enabling prediction of phenotype evolution from the epigenomic state^32,33^. This micro-/macro-state model is analogous to the thermodynamic situation where one can measure the energy of a single particle (microstate), but that measurement does not provide any information about the temperature of the system as a whole (macrostate). Like energy and temperature in thermodynamics, we have shown that the CML disease phenotype is encoded into a transcriptome macrostate that is defined by the ensemble of cell microstates.

Of note, we demonstrated that different PBMC cell types contributed varying amounts of information to the identification of CML development dynamics (Fig. 2C). This enabled us to develop a mechanistic model explaining how the BCR::ABL oncogene, while necessary for CML development, differentially affects each cell type’s transcriptome, and how these PsB cell types collectively reproduce the leukemia dynamics observed in bulk RNA data. Importantly, these observations applied to two different mouse models, one that recapitulated CP and the other BC. Of note, we observed that all the cell types had non-linear dynamics similar to the observed disease dynamics in the PBMCs. However, one intriguing result is that even monostable cell type dynamics when combined could produce tristable system-level dynamics as those observed from the PsB and bulk transcriptome analysis. This modeling result mathematically demonstrates how simple, linear dynamics within individual cell types can generate complex, non-linear phenotype dynamics. Furthermore, it provides a mechanistic and mathematical framework explaining how biological systems can exhibit emergent properties where the whole exceeds the sum of its parts.

In conclusion, our findings establish a novel conceptual framework for understanding how cellular information generate phenotypic states that define disease state-transitions, with wide ranging potential clinical applications in predicting disease development, transformation, and treatment response. By quantifying how single-cell transcriptional microstates comprise disease-defining macrostates, our framework suggests a path toward predictive modeling that could inform clinical management of CML by identifying patients at highest risk of progression or optimizing therapeutic strategies. Future application of these models in patient care may enable more accurate predictions of disease progression and treatment outcomes, facilitating timely therapeutic intervention.

## METHODS

### Mouse model and experimental design

The SCLtTA/BCR::ABL and Mir142^−/−^SCLtTA/BCR::ABL mice in B6 background were maintained on tetracycline (tet) water at 0.5 g/L. Tet withdrawal (tet-off) results in expression of BCR::ABL and generation of a CP CML-like disease in SCLtTA/BCR::ABL mice with a median survival of approximately 55 days and of a BC CML-like disease in Mir142^−/−^SCLtTA/BCR::ABL mice with a median survival of approximately 32 days, after induction of BCR::ABL. Blood (100μl) was collected from six to eight weeks old male (n=1) and female (n=2) SCLtTA/BCR::ABL or Mir142^−/−^SCLtTA/BCR::ABL mice weekly, i.e., before BCR::ABL induction (t = 0) and after induction up to 9 weeks (t = 1 to 9) or when the mouse developed leukemia and became moribund, whichever event occurred first. After red cell lysis, an aliquot of the PBMCs was stained with anti-mouse antibodies and analyzed for hematopoietic cell subpopulations and the remaining cells were subjected to scRNA-seq.

### Sequencing and data processing

Approximately 10,000 cells per murine sample were captured using a Chromium X (10X Genomics, Pleasanton, CA), with the Chromium Next GEM Single Cell 3’ Reagent Kit V3.1 (10X Genomics, Cat. PN-1000268). Prior to sequencing, library quality was assessed using a High Sensitivity DNA Chip (Agilent, Santa Clara, CA, Cat. 5067-4626) and quantified with the Qubit High Sensitivity DNA Assay Kit (Thermo,Fisher Scientific, Waltham, MA, Cat. Q32854). Sequencing was performed on a NovaSeq 6000 (Illumina, San Diego, CA) using the S4 Reagent Kit v1.5, 200 cycles (Part no 20028313). Illumina Real-Time Analysis (RTA3) v3.4.4 was used for base calling and quality scores and Illumina BCL Convert v3.9.3 to export FASTQ files. All protocols were performed according to the manufacturer’s instructions. Comprehensive library quality assessment was performed using v2.1.0 of the nf-core/scrnaseq pipeline^34^ and reads (including those from introns) counted using 10X Genomics Cellranger count v7.0.0 implemented therein. We used the Gencode M28 reference annotation and genome (https://www.gencodegenes.org/, downloaded 22 March 2022) augmented with a custom

BRC::ABL transcript (GenBank accession forthcoming) prepared using the default parameters and preprocessing described for Cellranger mkref. To perform quality control, each library was filtered to remove genes expressed in fewer than 3 cells, and cells with fewer than 200 genes expressed, with more than 20% mitochondrial transcripts, or with fewer than 5% ribosomal transcripts, then merged into a single v5 Seurat^35^ object for downstream analysis. Cell types were assigned with the default parameters of SingleR^36^ (v2.6.0) using reference expression profiles from the Immunological Genome Project, accessed through the celldex package (v0.1.16.0). Raw sequence reads and the Seurat object are available via the Gene Expression Omnibus (GSE296507).

### State-space exploration at the sc-level

Like the previously described bulk RNA-seq CML state-space^6^ which separated health from leukemic transcriptional state and defined continuous trajectories for our time-series samples, the scRNA-seq data was investigated for a similar CML state-space. Here the goal was to find a lower dimensional representation of the data that captured the progression of a sample from health to leukemia in either all cells or a subset of cell types. We expected an sc-level CML state-space to have similar properties such that the healthy (T_0_) cells and the leukemic (T_f_) cells would occupy distinct transcriptional states defined by a lower dimensional representation of the data. Different dimensionality reduction techniques were applied to the sc-level data using all cells and all samples as input. PCA was performed using the Seurat package on the log normalized mean centered gene expression^35^. Based on the elbow of the scree plot of the singular values, the first six PCs were assessed for cells occupying unique transcriptional state associated with the healthy (T_0_) and leukemic (T_f_) states (Fig. 1A; S1B,I). UMAP and two scRNA-seq analysis tools, Decipher and Mellon, were also used to try to identify a CML state-space (Fig. S1B,E,F)^8,9^. Mellon was used with default parameters to identify cell clusters that changed density over leukemia development but did not reveal any cell clusters that were uniquely associated with disease state (Fig. S1E). Because Decipher was able to construct a latent space where divergent AML trajectories between sorted cells from healthy and leukemic bone marrow samples^9^, we applied it to our more diverse PBMC CML samples. To train the Decipher model, healthy time points (T_0_) were used as controls and the leukemic time points (T_f_) were used as leukemic samples. Decipher was applied to all cells and each of the four cell types to try to identify cell type specific disease trajectories; however, none of the Decipher latent spaces constructed trajectories that described leukemic development in our CML mice.

Each cell type was also analyzed separately to determine whether an sc-level state-space could be identified (Fig. S1D). For each cell type, SVD was performed on the log-normalized mean centered expression of the cells from the healthy (T_0_) and leukemic (T_f_) time points to decrease computation time. The cells from the remaining time points were then projected into the resulting PCs and visualized to try to identify any PC or combination of PCs where the cells from the leukemic time points occupied a unique transcriptional state. To ensure that selecting only the terminal time points (T_0_ and T_f_) did not probit a leukemic trajectory of cell states from being detected, SVD was also performed on 500 randomly selected cells from each mouse so that all time points would be included (data not shown). This analysis was repeated for the BC CML mice.

### PsB state-space construction

The PsB data was constructed by summing gene counts within each mouse’s time point samples. For each sample, the counts were normalized to counts per million (CPM). This resulted in a sample by gene expression (n=29,513) matrix similar to bulk RNA-seq had been performed on each PBMC sample. SVD was performed on the mean-centered, log2 normalized PsB CPM which decomposes the gene expression matrix (*X* = *U*Σ*V*^*^) into: *U* which provides components of the data in decreasing total variance explained (i.e. the principal components [PC]), Σ which are the eigenvalues of each PC, and *V*^*^ which are the loading values for how the genes are combined to construct each PC (Fig. S1J). The top 6 PCs were visually inspected using scaled time to identify whether they encoded mouse trajectories that separated healthy (T_0_) from leukemic (T_f_) time points, and all PCs were correlated with the BCR::ABL expression in each PsB sample (Fig. 1SH; Table S2). The BCR::ABL expression was determined by aligning the scRNA-seq reads to the BCR::ABL fusion gene transcript, normalizing based on the total aligned reads per each cell, and then summing the cell level reads to get the PsB BCR::ABL expression.

### CML potential

The PsB state-space was compared to the bulk state-space by identifying healthy and leukemic reference, or ‘anchor’ points and then scaling the bulk state-space to span the PsB state-space. For the bulk state-space the healthy anchor point was the mean bulk state-space coordinate of all week 0 samples, and the leukemic anchor was the mean bulk state-space coordinate of final time point for all BCR::ABL induced mice that developed leukemia. For the PsB state-space, the healthy anchor point was the mean PsB state-space coordinate of all the week 0 samples, and the leukemia anchor point was the mean PsB state-space coordinate of final time point for the mice that developed leukemia. Using these anchor points, the bulk state-space coordinates of all samples and the location of the state-space critical points were translated into the PsB state-space (Fig. S1K). We observed that the bulk and PsB trajectories described similar dynamics in the combined state-space. Although the PsB mice developed leukemia quicker, this was expected since the mice used in this experiment were homozygous BCR::ABL knock in whereas the mice used in the bulk RNA-seq experiment were heterozygous for BCR::ABL and consequently would receive reduced oncogene expression upon induction. Given the similar dynamics and higher density of samples in the bulk state-space, we used the translated bulk state-space critical points and resulting potential for the PsB which was constructed as previously described^6^.

### SVD and projection ctPsB samples into the state-space

For each cell type, a cell type pseudobulk sample was constructed for each time point sample. The ctPsB was constructed using the same data and method as the PsB samples except only the cells of each cell type were included. For each cell type, the ctPsB data was mean-centered and log2 normalized before performing SVD (Fig. S1J). The top size principal components were investigated for scaled time and other descriptive variables to explain the top sources of variance in each cell type. The ctPsB CPM values were also used to project each ctPsB into the PsB state-space. First, the ctPsB data were mean centered using the means from the PsB data and then they were log2 normalized. To project the data into the PsB state-space, the PsB principal components of the ctPsB were calculated as *U*_*ctPsB*_ = *X*_*ctPsB*_*V*Σ^−1^, where *X*_*ctPsB*_ are the mean-centered log2 ctPsB expression matrix and *V* and Σ^−1^ are, respectively, the loading values and singular values from the PsB SVD results (Fig. S1J). For both the ctPsB SVD results (Fig. S2B) and the ctPsB projections (Fig. S2C), PC1, which corresponded to the PsB state-space, was plotted with the PsB state-space vs time for each cell type separately to observe the location and dynamics each ctPsB had in the PsB state-space. The critical points for each set of ctPsB trajectories and the resulting potentials were estimated by observing the ctPsB trajectory dynamics in the ctPsB state-spaces (Fig. S2B) and when projected into the PsB state-space (Fig. S2C). For the ctPsB projections, the location and the distance that each set of trajectories overlapped the PsB trajectories was summarized using the maximum and minimum values observed for each cell type to create a spanning vector (Fig. S2G).

### Cell type population changes

The number of each of the four categorized cell types identified in each sample was used to calculate the proportional representation of each cell type and plotted over scaled time (Fig. 2A; Table S3). The samples were then grouped by disease states, as defined by the critical points from the PsB state-space, and *propeller* from the *speckle* package in R^37^. A false discovery rate (FDR) p-value < 0.05 was used to determine significant changes in cell type populations.

### Fixed cell type computational simulation

The contribution of each cell type to the state-space trajectories was assessed using a computational simulation strategy. The simulation was performed separately for each cell type. For the cell type being simulated, all cells identified as that cell type were removed from each time point sample. Then, the gene expression for all the week 0 cells identified as that cell type were added to each time point sample (Fig. 2B). The result for each mouse was that one cell type was fixed to the week 0 cell count and transcriptional state while all other cell types underwent CML development. Next, the fixed simulation for each time point sample was aggregated into PsB data. The PsB data were then projected into the PsB state-space so that the fixed simulation trajectories for each cell type could be compared with the observed PsB trajectories for each mouse (Fig. 2C). The percent distance between each cell type’s fixed simulation and the observed PsB trajectory was calculated by first taking the difference between the observed and the fixed simulation PsB state-space coordinates for each time point sample. Those differences were then divided by the maximum distance spanned by each mouse’s observed trajectory which was the difference between the maximum and the minimum PsB state-space coordinate for each mouse. Finally, a fit line was drawn for the resulting percent differences of each cell type and plotted as a function of time (Fig. 2C middle). The difference in information contained in the trajectories of each fixed cell type simulation vs the observed trajectories was calculated using the Kullback-Leibler divergence which takes larger values when the simulated trajectories encode less of the observed trajectories (Fig. 2C right).

### Application of state-transition to BC samples

The blast crisis CML data were collected using an identical time-series experiment as the CP CML mice. The BC CML scRNA-seq samples were processed identically and all analyses were conducted as described in the CP CML mice. The PsB state-space was determined by performing SVD on the PsB gene expression matrix, which also identified PC1 as the PsB state-space for BC CML (Fig. S3D). Since the BC CML genotype differed from the CP CML genotype used in both the scRNA-seq and bulk RNA experiments, PsB state-space dynamics and resulting potential were determined using only the BC CML mice. By assessing the sample distribution in the PsB state-space using both histograms and kernel density, the BC CML state-space was determined to have two stable critical points and one unstable critical point (Fig. S3D). Using the same approach applied to the CP CML mice, the ctPsB state-spaces were identified to be encoded in PC1 for each cell type (Fig. S3F). When the ctPsB data were projected into the PsB state-space, the ctPsB trajectories were shown to have a different location and contribution to the PsB state-space (Fig. S3G). Using the BC CML PsB state-space, the fixed cell type simulation strategy was applied to the BC CML mice to quantify the information lost when each cell type was fixed (Fig. 2C).

### Human CD34+ BM samples and analysis

Cryopreserved human healthy pediatric CD34+ bone marrow samples were commercially purchased from STEMCELL Technologies (Milpitas, CA; Cat. 7000.2) and Lonza Bioscience (Walkersville, MD; Cat. 2M-101). Cryopreserved bone marrow aspirates from pediatric CML bone marrow patients at diagnosis were obtained from the Stanford Bass Center Tissue Bank. Lineage-/CD34+ cells were isolated by fluorescence-activated cell sorting for the 4 pediatric CML bone marrow samples. All samples were processed by collaborating with the Stanford Functional Genomics Facility. The Chromium Next GEM Single Cell 3’ Reagent Kit (10x Genomics, Pleasanton, CA; cat. PN-1000268) was used for single cell capture and library preparation. Captured cell counts varied across samples, as healthy samples contained a median of ∼6,200 cells (range: 3,200–8,300), while CML samples contained a median of ∼6,100 cells (range: 800– 11,500). The libraries were sequenced on a NovaSeq 6000 (Illumina, San Diego, CA). Reads were counted using 10x Genomics Cellranger count v6.0.0 with the 10x GRCh38-2020-A reference transcriptome. Libraries were then aggregated with Cellranger aggr v6.0.0 prior to being combined into a single v4 Seurat object. To perform quality control, the Seurat object was filtered to remove genes expressed in fewer than 3 cells, and remove cells with fewer than 200 genes expressed, greater than 8000 genes expressed, or with more than 10% mitochondrial transcripts. Cell types were assigned with the default parameters of SingleR^36^ (v2.6.0) using reference expression profiles from the Human Primary Cell Atlas, accessed through the celldex package (v0.1.16.0; Fig S4A). Raw sequence reads and the Seurat object are available via the Gene Expression Omnibus (Accession number forthcoming). PCA and UMAP was performed for all cells and for cells of each cell type as previously described to try to identify emergent cell states (Fig 3A-B;S4B-C). For each sample, a PsB sample was constructed and SVD was performed on all PsB samples to identify the CML state-space in PC2. Fixed cell type simulations were performed as previous described, except to accommodate the unpaired samples in this data set, each CML sample (n=4) was paired with a healthy sample (n=3) and all possible pairings of CML and healthy samples were performed by performing the fixed cell type simulation on each of four possible groupings (Fig. S4D). The information lost from each simulation was calculated by comparing the full and simulated trajectories for each sample pairing using Kullback-Leibler divergence as previously described and the mean divergence across the four groupings was reported (Fig. S4D).

### PsB and ctPsB model

In the PsB model, only the transcriptome (*U*) is a linear combination of the cell types with their weights (*w*_*i*_) where each cell type evolves independently and was described by:

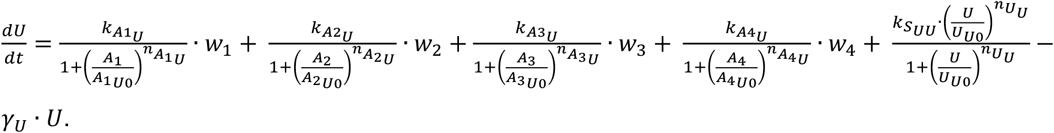

Here, *w*_1_, *w*_2_, *w*_3_, *w*_4_ are the weightages of the four cell types whose value decides the contribution of each cell type to the PsB transcriptome. The individual cell types evolve independently according to the equations:

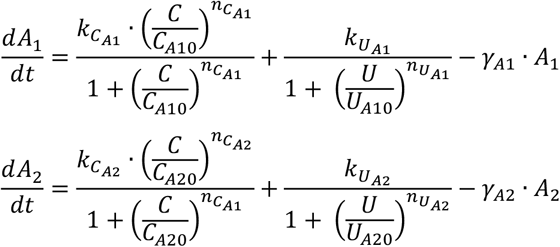

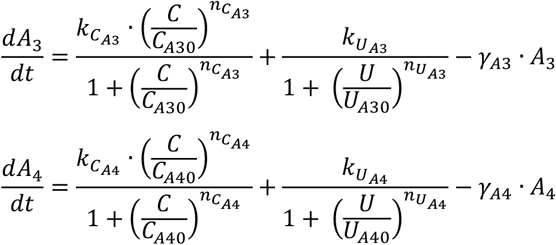

This model is represented in Figure S5B with *w*_*i*_ terms shown in the representation in the upper right. The individual cell type were produced from this model by setting the value of *w*_*i*_ to 0 or 1. For cell type in the ctPsB models:

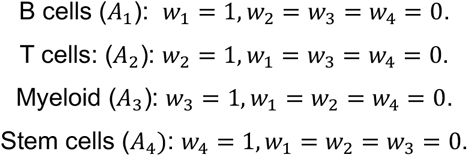

For the PsB model we set *w*_1_ = *w*_2_ = *w*_3_ = *w*_4_ = 1 which is represented in upper left and bottom of Figure S5B. In the ODEs, 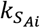 and 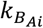 represent the maximum expression of *A*_*i*_ due to *C* and *U*, respectively. *C*_*Ai*0_ represent the threshold value of *C* at the half-maximum of 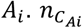 and 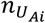 represent the Hill coefficient on *A*_*i*_ due to *C* and *U*, respectively. The Hill coefficient on U due to *A*_*i*_ and *U* is given by 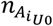 and 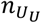, respectively. 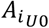 and *U*_*U*0_ are the threshold of *A*_*i*_ and *U*, respectively at the half-maximum value of *U*.

For a given model, the general form of the ODE would be

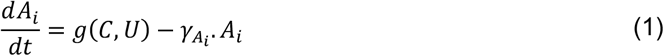

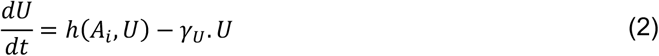

The steady state equation of *A*_*i*_ is written as:

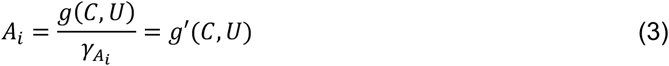

Substituting the steady state equation of *A*_*i*_ in *U* using a transfer function, we get

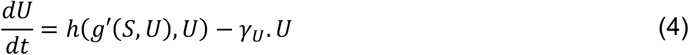

Integrating the negative of Equation 4 from 0 to *U* gives the potential function *U*(*C, U*)

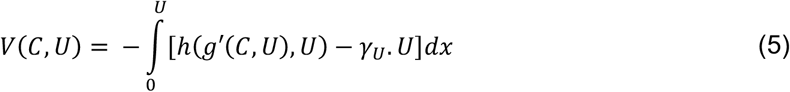

Following the method introduced by Dey and Barik^11^, using Equation 5, we first generated the different bifurcations for the ctPsB and PsB models. Using these bifurcations with the parameter values indicated in Table S4, we then generated the potential landscapes (Fig. 4B).

## Data and code availability

All scRNA-seq raw sequence reads and the Seurat object are available via the Gene Expression Omnibus for the mouse CP and BC CML (GSE296507) and human (Accession number forthcoming) experiments. Code to perform all analyses, modeling, and to generate all figures are available at https://github.com/cohmathonc/CML.BC.scRNA-manuscript.

## Acknowledgements

Research reported in this publication was supported in part by Robert and Lynda Altman Family Foundation and included work supported by the City of Hope Integrative Genomics Core, High Performance Research Computing Center and Biostatistics and Mathematical Oncology shared resources supported by the National Cancer Institute of the National Institutes of Health under grant numbers P30CA033572, U01CA250046, U01CA293853; Stanford Maternal and Child Health Institute, Lurie Children’s Hospital/Northwestern University (K.M.S); American Society of Hematology (T.Y); Alex’s Lemonade Stand Foundation (R.S.). The content is solely the responsibility of the authors and does not necessarily represent the official views of the National Institutes of Health.

## Author contributions

Conceptualization, GM, BZ, RCR, and Y-HK; Methodology, RCR, DEF, ALM, AD, SB; Investigation, DZ and BZ; Formal Analysis, DEF, RCR, ZC, JI, AD, and DO. Writing – Original Draft, DEF, RCR, KMS and GM; Writing – Reviewing & Editing, all authors; Supervision, RCR and GM; Funding Acquisition, GM, RCR, Y-HK.

## Supplementary Figures

**Figure S1:**
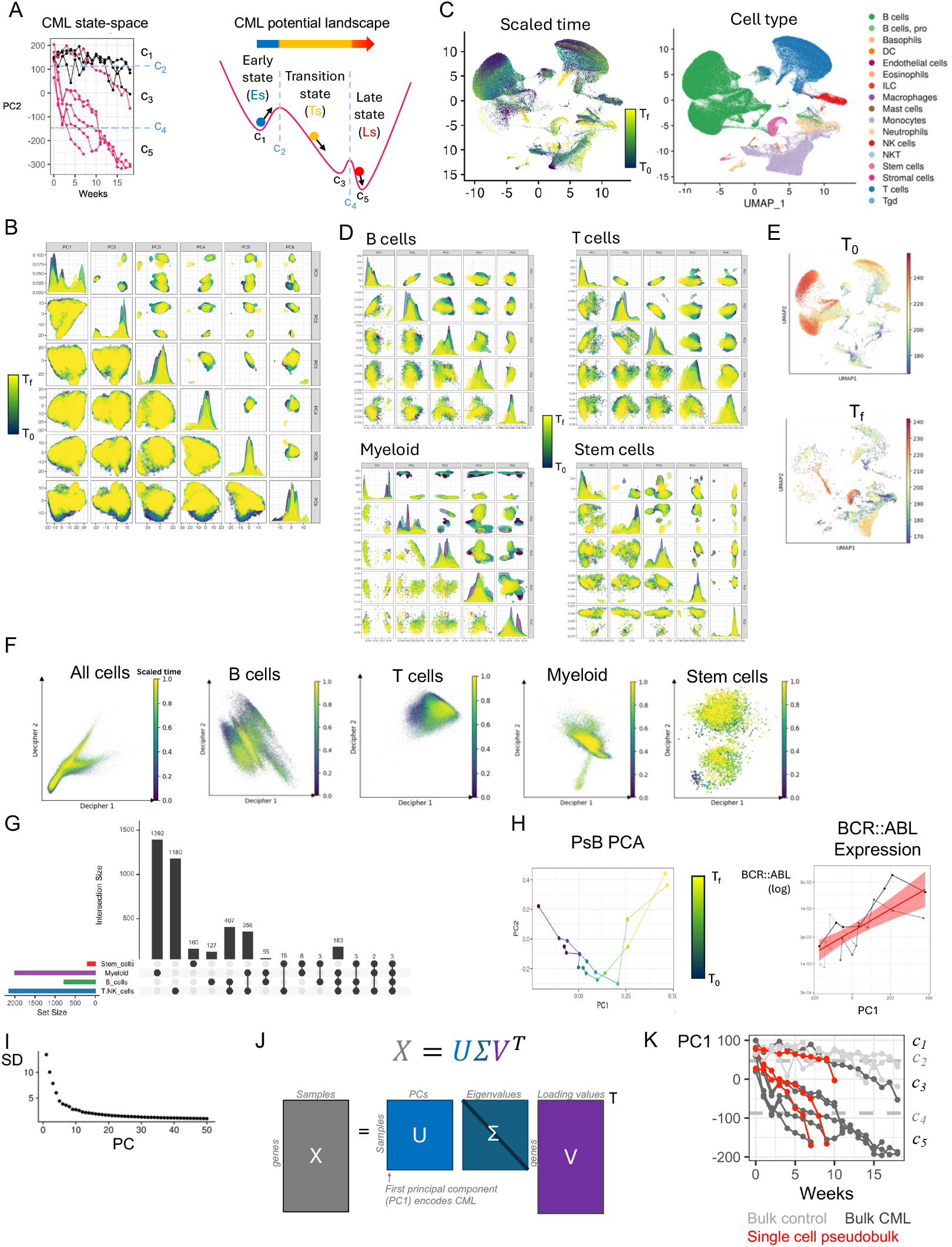
**A**) Mouse trajectories in the CML state-space constructed from the time-series bulk RNA-seq data (*left*)^6^. See Frankhouser et al. 2024 for full details on state-space construction, critical point identification, and potential landscape determination. The representative three-well potential describes how each mouse was modeled as undergoing Brownian motion in a potential energy landscape (*right*). The critical points of the potential were used to define the phenotypic disease states (Es, Ts, Ls). **B**) PCA was performed on the mean-centered expression of all genes using all cells from the time -series scRNA-seq data from the CP CML mice. For the mice that developed CML, they progressed at different rates so each mouse’s week 0 (T_0_) and their final time point (T_f_) were scaled range between 0 and 1 so that their CML trajectories were aligned. The scaled time was used to try to identify a state-transition at the sc-level in PC1-5 where the elbow of the scree plot occurred (Fig. S1I). **C**) UMAP was also performed on all cells from all CP CML samples to try to identify a sc-level state-transitions using scaled time. Labeling each cell by the cell type shows that UMAP primarily separates different cell populations of the PBMCs. **D**) For each cell type category, SVD was performed on the mean-centered gene expression and PCs 1-5 were colored by the scaled time of the leukemic mice to identify a leukemia state-transition within cell type at the sc-level. **E**) The cell state density tool Mellon was used to try to identify sc-level leukemic state-transitions^8^. **F**) Decipher, a tool for constructing a latent space from control and perturbed cell populations to identify trajectories of derailed cells, was also used on all cells and on each cell type separately^9^. After building the Decipher space using the cells from the healthy time points vs the post induction time points, the resulting latent spaces were colored using the samples scaled time. **G**) Upset plot of the DEGs identified by comparing the cells from healthy (T_0_) vs leukemic (T_f_) time point samples from the leukemic mice for each cell type. **H**) The mouse pseudobulk (PsB) trajectories are shown in PC1 vs PC2 (*left*). PC1 was identified as the PsB state-space as it was the only PC that showed concordance with the expression of the BCR::ABL leukemia oncogene (*right*). **I**) Scree plot of the standard deviation (SD) explained by the PCs that result from performing PCA on the mean-centered gene expression of the sc-level scRNA-seq data. **J**) Singular value decomposition (SVD) equation and a representation of how it was applied to the PsB gene expression matrix.

**Figure S2:**
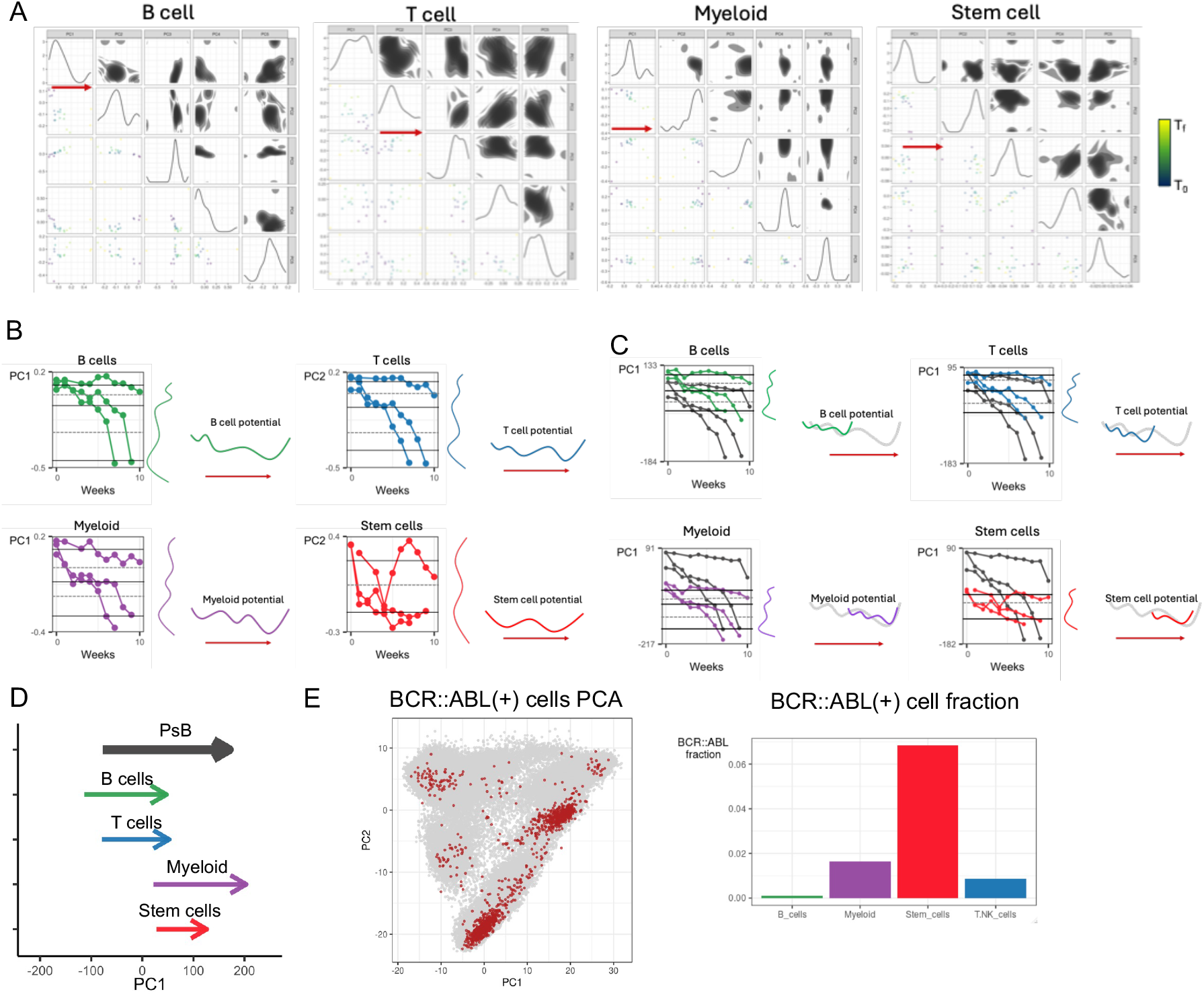
**A**) SVD was performed on each cell type pseudobulk (ctPsB) gene expression matrix; the first five PCs were plotted usind the scaled time colors, and the PC that encoded the ctPsB state-space was indicated (*red arrows*). **B**) For each ctPsB state-space, the trajectories were used to estimate the state-space dynamics for each cell type. The stable critical points (*black lines*) were drawn based on how many stable states were observed, and the unstable critical points (*grey dashed lines*) were drawn based on where the trajectories showed the highest velocity. Using these critical points, an estimated potential was constructed to illustrate the dynamics of each cell type during leukemia progression (*red arrow*). **C**) The ctPsB data for each cell type (*colored trajectories*) were projected into the PsB state-space (PsB trajectories in shown in black) by multiplying the ctPsB gene expression by the right singular values from the PsB SVD result (Fig. S1J). For each ctPsB projection, the stable critical points (*black lines*) were drawn based on how many stable states were observed, and the unstable critical points (*grey dashed lines*) were drawn based on where the trajectories showed the highest velocity. Using these critical points, an estimated potential was constructed to illustrate the dynamics of each cell type during leukemia progression (*red arrow*). The ctPsB potentials were shown with respect to the PsB potential (*grey*) constructed using the same critical point estimation approach. **D**) The extent and location of each ctPsB projected trajectories and the PsB trajectories were each summarized using the minimum and maximum PsB state-space coordinate. **E**) Cells with BCR::ABL expression [BCR::ABL(+)] were indicated in the PCA representation of all cells from all time points (*left*). The bar plot (*right*) shows the fraction of BCR::ABL(+) cells in each cell type.

**Figure S3:**
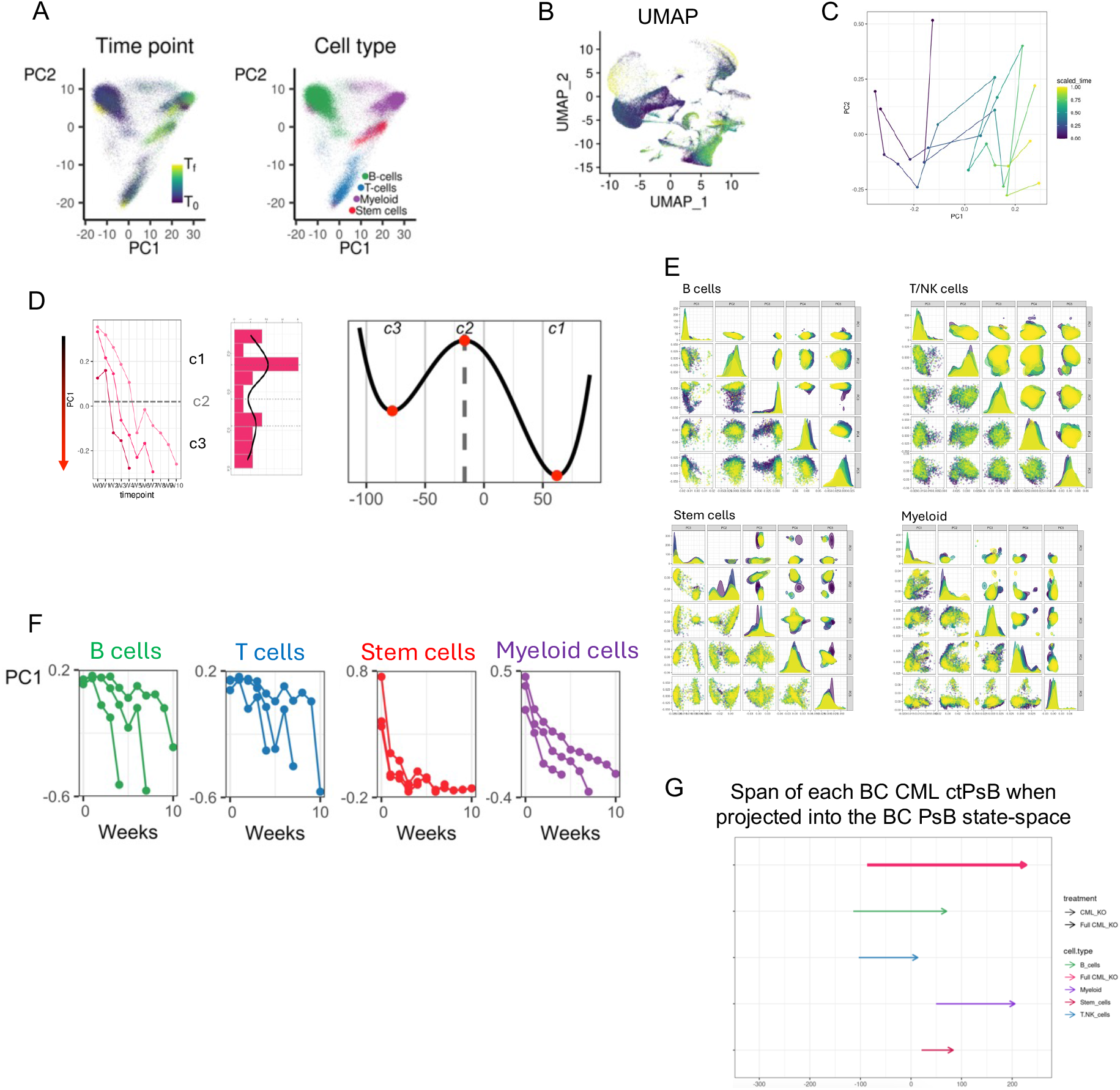
**A**) Using all cells and all time point of the BC CML mouse experiment, PCA was performed on the mean-centered expression of all genes. The first two PCs were plotted for each cell, and the cells were colored by the sample’s scaled time points (*left*) and cell type (*right*). **B**) UMAP was also performed on all cells, and the cells were colored by their samples scaled time. **C**) SVD was performed on the BC CML PsB gene expression matrix, and the mouse trajectories in the first two PCs were shown and colored by the scaled time. PC1 encoded the BC CML PsB state-space. **D**) The trajectories of each BC CML mouse are shown in the PsB state-space (*left*). The sample density in the PsB state-space was fit using kernel density estimate to determine the number of steady states and the location of the critical points (*middle*). The critical points were used to construct a potential function that describes the PsB BC CML dynamics. **E**) SVD was performed on the sc-level gene expression separately for each cell type, and the sample’s scaled time was shown used to color each cell in the first five PCs of each cell type. **F**) SVD was performed on the ctPsB data for each cell type and the mouse ctPsB trajectories were shown for PC1 which was identified as the ctPsB state-space for each cell type. **G**) The ctPsB gene expression data were projected into the PsB state-space for each cell type. The extent and location of the ctPsB trajectories and the PsB trajectories were summarized by taking the minimum and maximum PsB state-space coordinate for each set of trajectories.

**Figure S4:**
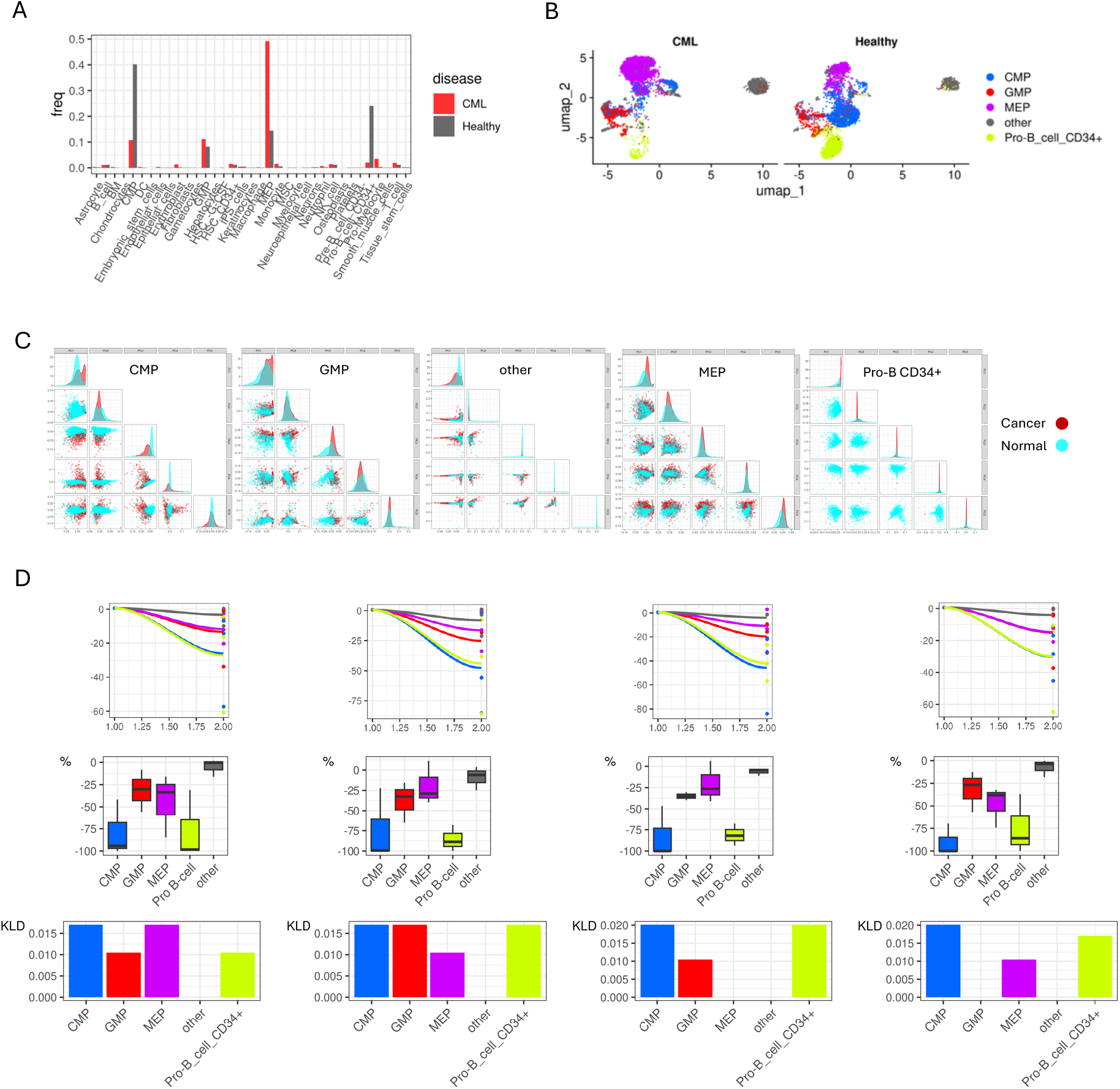
**A**) From the human CD34+ BM scRNA-seq data, cell types were labeled and summarized as the fraction of total cells in each disease group. For the purposes of analysis, all cell types with less 5% in both healthy and CML disease groups were categorized as “other”. **B**) UMAP was performed on all cells, and the cells were colored by cell type. **C**) SVD was performed separately on each cell type, and the top five PCs were used to try to identify transcriptional cell states unique to the CML samples. **D**) To quantify how each cell type contributed to CML development, the unpaired samples were treated as healthy-CML pairs by matching each of the three healthy samples to one of the CML samples to create four different healthy-CML pair combinations. All possible combinations of sample pairings were used in the fixed simulations, and for each pairing and each cell type, the CML sample’s cells were replaced with the paired healthy sample’s cells to simulate what would happen if each cell type was fixed in a healthy state. The plots here quantify how fixing each cell type moved the location of the CML sample in the state-space toward the healthy state (top row) and the percent of the total distance observed for each healthy-CML pair when the cell types were not fixed at the healthy state (middle row). The information lost by fixing each cell type was computed using Kullbach-Leibler divergence (KLD; bottom row).

**Figure S5:**
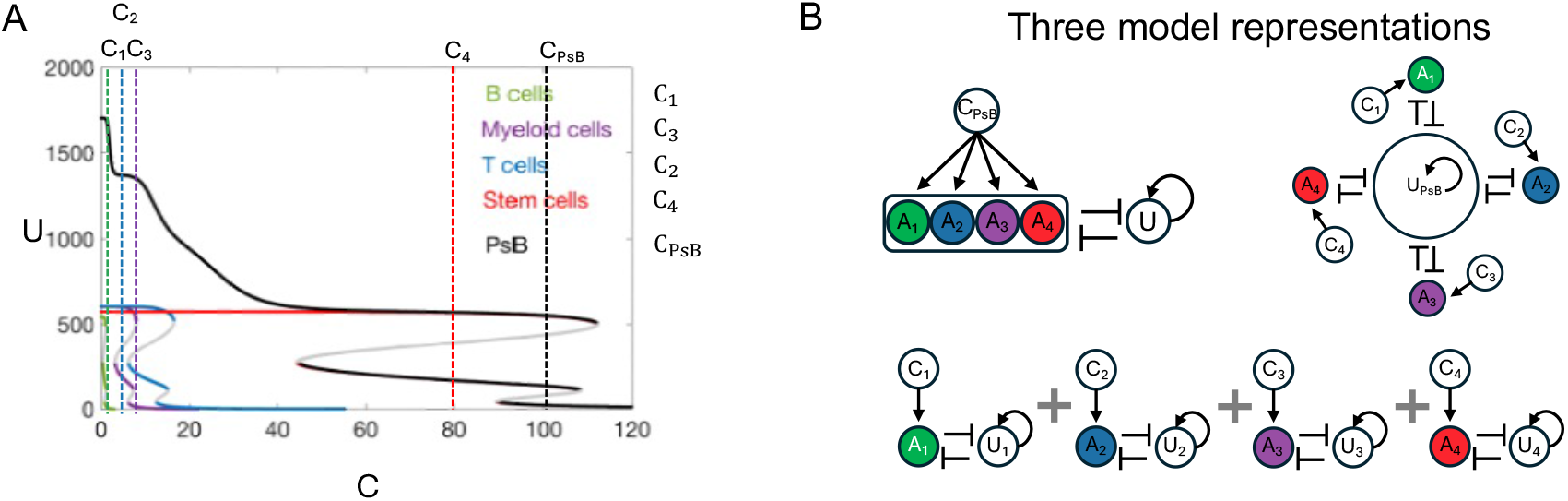
**A**) The bifurcation diagram that produces the ctPsB and PsB potentials shown in Figure 3B. The potentials are determined by setting BCR::ABL signal levels (*C*_*ct*_) in these bifurcations as *C*_1_ = 1; *C*_2_ = 7; *C*_3_ = 14; *C*_4_ = 80, which results in a PsB BCR::ABL signal of *C*_*PsB*_ = 102. **B**) Three representations of the model that related the ctPsB dynamics to the PsB dynamics.

## Supplementary Tables

**Table S1: Cell type counts and categories**

**Table S2: PsB state-space loading values and correlation with BCR::ABL**

**Table S3: Cell type proportion testing table**

**Table S4: Model parameter values**

