## Supplementary material for "Longitudinal single cell RNA-sequencing reveals evolution of micro- and macro-states in chronic myeloid leukemia"

**Supplementary Figures:**

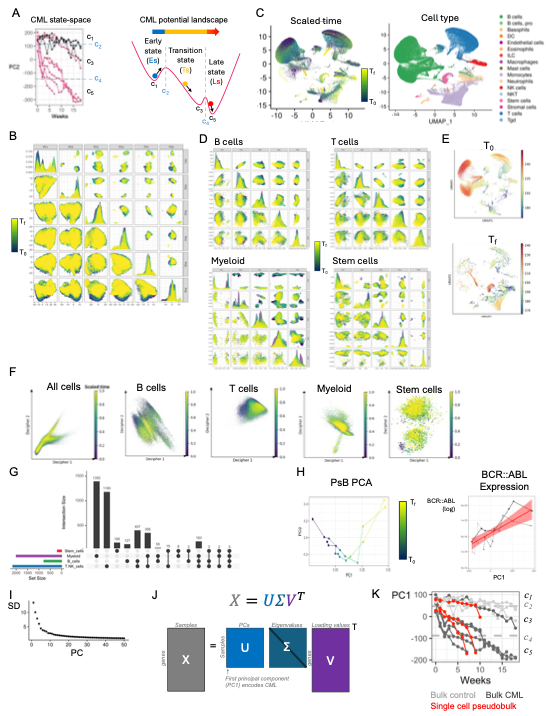

**Figure S1**:

**A**) Mouse trajectories in the CML state-space constructed from the time-series bulk RNA-seq data (*left*)^4^. See Frankhouser et al. 2024 for full details on state-space construction, critical point identification, and potential landscape determination. The representative three-well potential describes how each mouse was modeled as undergoing Brownian motion in a potential energy landscape (*right*). The critical points of the potential were used to define the phenotypic disease states (Es, Ts, Ls). **B**) PCA was performed on the mean-centered expression of all genes using all cells from the time -series scRNA-seq data from the CP CML mice. For the mice that developed CML, they progressed at different rates so each mouse’s week 0 (T_0_) and their final time point (T_f_) were scaled range between 0 and 1 so that their CML trajectories were aligned. The scaled time was used to try to identify a state-transition at the sc-level in PC1-5 where the elbow of the scree plot occurred (Fig. S1I). **C**) UMAP was also performed on all cells from all CP CML samples to try to identify a sc-level state-transitions using scaled time. Labeling each cell by the cell type shows that UMAP primarily separates different cell populations of the PBMCs. **D**) For each cell type category, SVD was performed on the mean-centered gene expression and PCs 1-5 were colored by the scaled time of the leukemic mice to identify a leukemia state-transition within cell type at the sc-level. **E**) The cell state density tool Mellon was used to try to identify sc-level leukemic state-transitions^6^. **F**) Decipher, a tool for constructing a latent space from control and perturbed cell populations to identify trajectories of derailed cells, was also used on all cells and on each cell type separately^7^. After building the Decipher space using the cells from the healthy time points vs the post induction time points, the resulting latent spaces were colored using the samples scaled time. **G**) Upset plot of the DEGs identified by comparing the cells from healthy (T_0_) vs leukemic (T_f_) time point samples from the leukemic mice for each cell type. **H**) The mouse pseudobulk (PsB) trajectories are shown in PC1 vs PC2 (*left*). PC1 was identified as the PsB state-space as it was the only PC that showed concordance with the expression of the BCR::ABL leukemia oncogene (*right*). **I**) Scree plot of the standard deviation (SD) explained by the PCs that result from performing PCA on the mean-centered gene expression of the sc-level scRNA-seq data. **J**) Singular value decomposition (SVD) equation and a representation of how it was applied to the PsB gene expression matrix.

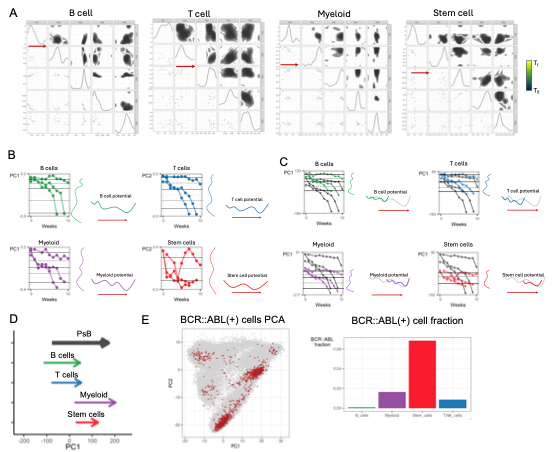

**Figure S2:**

**A**) SVD was performed on each cell type pseudobulk (ctPsB) gene expression matrix; the first five PCs were plotted usind the scaled time colors, and the PC that encoded the ctPsB state-space was indicated (*red arrows*). **B**) For each ctPsB state-space, the trajectories were used to estimate the state-space dynamics for each cell type. The stable critical points (*black lines*) were drawn based on how many stable states were observed, and the unstable critical points (*grey dashed lines*) were drawn based on where the trajectories showed the highest velocity. Using these critical points, an estimated potential was constructed to illustrate the dynamics of each cell type during leukemia progression (*red arrow*). **C**) The ctPsB data for each cell type (*colored trajectories*) were projected into the PsB state-space (PsB trajectories in shown in black) by multiplying the ctPsB gene expression by the right singular values from the PsB SVD result (Fig. S1J). For each ctPsB projection, the stable critical points (*black lines*) were drawn based on how many stable states were observed, and the unstable critical points (*grey dashed lines*) were drawn based on where the trajectories showed the highest velocity. Using these critical points, an estimated potential was constructed to illustrate the dynamics of each cell type during leukemia progression (*red arrow*). The ctPsB potentials were shown with respect to the PsB potential (*grey*) constructed using the same critical point estimation approach. **D**) The extent and location of each ctPsB projected trajectories and the PsB trajectories were each summarized using the minimum and maximum PsB state-space coordinate. **E**) Cells with BCR::ABL expression [BCR::ABL(+)] were indicated in the PCA representation of all cells from all time points (*left*). The bar plot (*right*) shows the fraction of BCR::ABL(+) cells in each cell type.

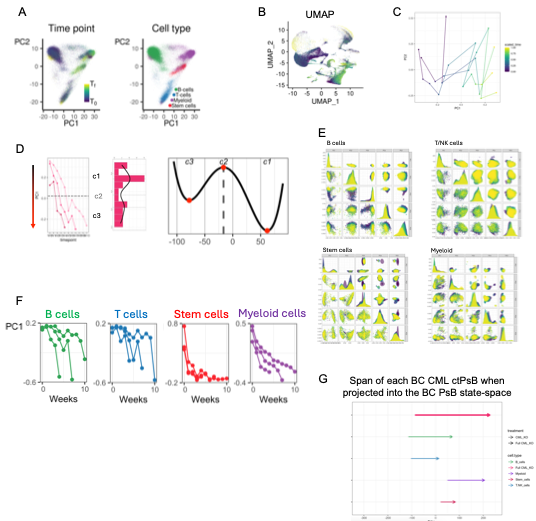

**Figure S3:**

**A**) Using all cells and all time point of the BC CML mouse experiment, PCA was performed on the mean-centered expression of all genes. The first two PCs were plotted for each cell, and the cells were colored by the sample’s scaled time points (*left*) and cell type (*right*). **B**) UMAP was also performed on all cells, and the cells were colored by their samples scaled time. **C**) SVD was performed on the BC CML PsB gene expression matrix, and the mouse trajectories in the first two PCs were shown and colored by the scaled time. PC1 encoded the BC CML PsB state-space. **D**) The trajectories of each BC CML mouse are shown in the PsB state-space (*left*). The sample density in the PsB state-space was fit using kernel density estimate to determine the number of steady states and the location of the critical points (*middle*). The critical points were used to construct a potential function that describes the PsB BC CML dynamics. **E**) SVD was performed on the sc-level gene expression separately for each cell type, and the sample’s scaled time was shown used to color each cell in the first five PCs of each cell type. **F**) SVD was performed on the ctPsB data for each cell type and the mouse ctPsB trajectories were shown for PC1 which was identified as the ctPsB state-space for each cell type. **G**) The ctPsB gene expression data were projected into the PsB state-space for each cell type. The extent and location of the ctPsB trajectories and the PsB trajectories were summarized by taking the minimum and maximum PsB state-space coordinate for each set of trajectories.

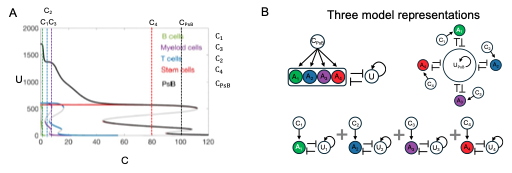

**Figure S4:**

**A**) The bifurcation diagram that produces the ctPsB and PsB potentials shown in Figure 3B. The potentials are determined by setting BCR::ABL signal levels ($C_{ct}$) in these bifurcations as $C_{1}=1;C_{2}=7;C_{3}=14;C_{4}=80$, which results in a PsB BCR::ABL signal of $C_{PsB}=102$. **B**) Three representations of the model that related the ctPsB dynamics to the PsB dynamics.

**Supplementary Tables:**

**Table S1: Cell type counts and categories**

**Table S2: PsB state-space loading values and correlation with BCR::ABL**

**Table S3: Cell type proportion testing table**

**Table S4: Model parameter values**
