## Supplementary material for "Longitudinal single cell RNA-sequencing reveals evolution of micro- and macro-states in chronic myeloid leukemia": Table S4

Parameter values for the model

| # | Parameters | Description | For ctPsB and PsB tristable | For ctPsB bi- and mono-stable and PsB tristable |
| --- | --- | --- | --- | --- |
| 1 | $k_{C_{A1}}$ | Maximum expression rate of $A_{1}$ due to $C$ | 200 | 1200 |
| 2 | $k_{C_{A2}}$ | Maximum expression rate of $A_{2}$ due to $C$ | 500 | 200 |
| 3 | $k_{C_{A3}}$ | Maximum expression rate of $A_{3}$ due to $C$ | 550 | 100 |
| 4 | $k_{C_{A4}}$ | Maximum expression rate of $A_{4}$ due to $C$ | 600 | 150 |
| 5 | $C_{A10}$ | Threshold of $C$ at half-maximum $A_{1}$ | 40 | 90 |
| 6 | $C_{A20}$ | Threshold of $C$ at half-maximum $A_{2}$ | 100 | 90 |
| 7 | $C_{A30}$ | Threshold of $C$ at half-maximum $A_{3}$ | 120 | 85 |
| 8 | $C_{A40}$ | Threshold of $C$ at half-maximum $A_{4}$ | 300 | 1000 |
| 9 | $k_{U_{A1}}$ | Maximum expression rate of $A_{1}$ due to $U$ | 25 | 25 |
| 10 | $k_{U_{A2}}$ | Maximum expression rate of $A_{2}$ due to $U$ | 22 | 25 |
| 11 | $k_{U_{A3}}$ | Maximum expression rate of $A_{3}$ due to $U$ | 25 | 25 |
| 12 | $k_{U_{A4}}$ | Maximum expression rate of $A_{4}$ due to $U$ | 28 | 25 |
| 13 | $U_{A10}$ | Threshold of $U$ at half-maximum $A_{1}$ | 400 | 400 |
| 14 | $U_{A20}$ | Threshold of $U$ at half-maximum $A_{2}$ | 400 | 400 |
| 15 | $U_{A30}$ | Threshold of $U$ at half-maximum $A_{3}$ | 400 | 400 |
| 16 | $U_{A40}$ | Threshold of $U$ at half-maximum $A_{4}$ | 400 | 400 |
| 17 | $k_{A1_{U}}$ | Maximum expression rate of $U$ due to $A_{1}$ | 40 | 55 |
| 18 | $k_{A2_{U}}$ | Maximum expression rate of $U$ due to $A_{2}$ | 45 | 65 |
| 19 | $k_{A3_{U}}$ | Maximum expression rate of $U$ due to $A_{3}$ | 48 | 65 |
| 20 | $k_{A4_{U}}$ | Maximum expression rate of $U$ due to $A_{4}$ | 45 | 60 |
| 21 | $k_{U_{U}}$ | Maximum expression rate of $U$ due to self-activation | 36.6837 | 36.6837 |
| 22 | $U_{U_{0}}$ | Threshold of $U$ at half-maximum $U$ (self-activation) | 125 | 125 |
| 23 | $A_{1_{U0}}$ | Threshold of $A_{1}$ at half-maximum $U$ | 125 | 150 |
| 24 | $A_{2_{U0}}$ | Threshold of $A_{2}$ at half-maximum $U$ | 130 | 150 |
| 25 | $A_{3_{U0}}$ | Threshold of $A_{3}$ at half-maximum $U$ | 150 | 150 |
| 26 | $A_{4_{U0}}$ | Threshold of $A_{4}$ at half-maximum $U$ | 150 | 150 |
| 27 | $n_{C_{A1}}$ | Hill coefficient of $C$ on $A_{1}$ | 1 | 1 |
| 28 | $n_{C_{A2}}$ | Hill coefficient of $C$ on $A_{2}$ | 1 | 1 |
| 29 | $n_{C_{A3}}$ | Hill coefficient of $C$ on $A_{3}$ | 1 | 1 |
| 30 | $n_{C_{A4}}$ | Hill coefficient of $C$ on $A_{4}$ | 1 | 1 |
| 31 | $n_{U_{A1}}$ | Hill coefficient of $U$ on $A_{1}$ | 6 | 6 |
| 32 | $n_{U_{A2}}$ | Hill coefficient of $U$ on $A_{2}$ | 6 | 6 |
| 33 | $n_{U_{A3}}$ | Hill coefficient of $U$ on $A_{3}$ | 6 | 6 |
| 34 | $n_{U_{A4}}$ | Hill coefficient of $U$ on $A_{4}$ | 6 | 6 |
| 35 | $n_{{A_{1}}_{U}}$ | Hill coefficient of $A_{1}$ on $U$ | 7 | 7 |
| 36 | $n_{{A_{2}}_{U}}$ | Hill coefficient of $A_{2}$ on $U$ | 7 | 7 |
| 37 | $n_{{A_{3}}_{U}}$ | Hill coefficient of $A_{3}$ on $U$ | 7 | 7 |
| 38 | $n_{{A_{4}}_{U}}$ | Hill coefficient of $A_{4}$ on $U$ | 7 | 7 |
| 39 | $n_{U_{U}}$ | Hill coefficient of $U$ on $U$ (self-activation) | 3 | 2 |
| 40 | $\gamma_{A_{1}}$ | Degradation rate of $A_{1}$ | 0.1653 | 0.1653 |
| 41 | $\gamma_{A_{2}}$ | Degradation rate of $A_{2}$ | 0.1653 | 0.1653 |
| 42 | $\gamma_{A_{3}}$ | Degradation rate of $A_{3}$ | 0.1653 | 0.1653 |
| 43 | $\gamma_{A_{4}}$ | Degradation rate of $A_{4}$ | 0.1653 | 0.1653 |
| 44 | $\gamma_{U}$ | Degradation rate of $U$ | 0.1653 | 0.1653 |
| 45 | $w_{1}$ | weightage of $A_{1}$ on $U$ | For $A_{1}$ only ctPsB, set  $w_{1}=1, w_{2}=w_{3}=w_{4}=0$ | |
| 46 | $w_{2}$ | weightage of $A_{2}$ on $U$ | For $A_{2}$ only ctPsB, set  $w_{2}=1, w_{1}=w_{3}=w_{4}=0$ | |
| 47 | $w_{3}$ | weightage of $A_{3}$ on $U$ | For $A_{3}$ only ctPsB, set  $w_{3}=1, w_{1}=w_{2}=w_{4}=0$ | |
| 48 | $w_{4}$ | weightage of $A_{4}$ on $U$ | For $A_{4}$ only ctPsB, set  $w_{4}=1, w_{2}=w_{3}=w_{1}=0$ | |
| 49 |  |  | For PsB, set  $w_{1}=w_{2}=w_{3}=w_{4}=1$ | |
